# Population-specific drivers of reproductive phenology in a widespread large carnivore, the gray wolf

**DOI:** 10.64898/2026.05.25.727694

**Authors:** Lauren M. Hennelly, Shivam Shrotriya, Shaheer Khan, Alireza Mohammadi, Mohammad Farhadinia, Luis Llaneza, Joseph Bump, Austin Homkes, Steve Windels, Mohammad Zafarul Islam, Yadvendradev V. Jhala, José Vicente López-Bao, Gretchen Roffler, Mihir Godbole, Indrajeet Ghorpade, Pierre Comizzoli, Nucharin Songsasen, Håkan Sand, Camilla Wikenros, Peter Wabakken, Barbara Zimmermann, Peter Mahoney, Dan Stahler, Erin Stahler, Matthew Metz, Kira Cassidy, Thomas Gable, Bilal Habib

## Abstract

Animals often rely on cues from the environment to time reproduction with optimal conditions. However, for most species, which environmental cues are used and how their use varies across a species’ distribution remains poorly understood. The gray wolf (Canis lupus) is found across a broad latitudinal range (12°N to 83°N) and diverse natural habitats, making them an ideal study species to explore these questions. Using 1,261 estimated birth dates of wild and captive wolves across their global range, we investigated how environmental variables influence reproductive phenology in this widespread large carnivore. Birth dates for wild wolves ranged from November 22^nd^ for Indian wolves (18°N) to June 15^th^ for Arctic wolves (80°N). The best predictor of timing of birth was increasing daylength during the mating season, followed by precipitation variability during the year. The timing of birth in captive wolves was highly similar to wild wolves at same latitudes, suggesting they use similar environmental cues to time reproduction. Near the equator where photoperiod is more stable, wolves remained seasonal and likely use climatic cues to time reproduction. Our work reveals populations may use different environmental cues to time reproduction, a characteristic that likely facilitates adaptation to diverse environments.

## Introduction

Timing of birth is a key life history trait that impacts reproductive success and fitness (<u>Bronson 1985</u>). Its timing is a result of trade-offs imposed by the environment and an organism’s physiology. In response to these trade-offs, animals have evolved strategies to precisely time their reproduction to coincide with optimal climatic and foraging conditions. These strategies often rely on environmental cues, which can be fixed in the environment, such as photoperiod, or more variable, such as temperature and rainfall (<u>Bradshaw and Holzapfel 2007</u>). Understanding the environmental cues that influence reproductive timing has recently regained interest, particularly for predicting how animals will respond to climate change (<u>Bronson et al. 2009</u>, <u>Walker et al. 2019</u>, Abrahms et al. 2022).

For animals in high latitudes, photoperiod is the most reliable indicator for when annual peaks of resource availability occur (<u>Bradshaw and Holzapfel 2007</u>). Photoperiod is reliable because it is fixed in the environment; its variation is directly due to Earth’s tilt and does not vary from year-to-year. Due to its consistency and direct link to resource availability, animals have evolved multiple ways to perceive photoperiod and use it to regulate reproductive timing (<u>Nakane and Yoshimura 2014</u>). As latitude decreases towards the equator, photoperiod becomes a less reliable cue to time reproduction. This is partly because daylength varies little over the year at lower latitudes and resource availability is primarily driven by rainfall and temperature, rather than photoperiod. As a result, animals in lower latitudes often rely on climatic cues, such as precipitation and temperature, to time reproduction (<u>Ogutu et al. 2015</u>, <u>Bergallo and Magnusson 1999</u>).

Despite being a central component of life history, we know little about reproductive biology of most animals – let alone the roles of environmental cues on reproductive timing (<u>Wildt et al. 2003</u>). Controlled experiments altering daylength in captive animals have shown photoperiod to be a major determinant of reproductive activity for some species, but not for others near the equator (Gower et al. 1992, <u>McGukin and Blacksaw 1992</u>, <u>Pevet 1988</u>, <u>Demas and Nelson 1998</u>, <u>Trillmich 2000</u>, O’Brien et al. 1993, <u>Heideman and Bronson 1990</u>). Experimental approaches have also shown temperature can interact with photoperiodic cues to modulate reproductive timing (<u>Rosmalen et al. 2022</u>). However, these experimental studies are limited to small, short-lived mammals, which are hypothesized to be more opportunistic breeders and less photoperiodic than large, longer-lived mammals (Bronson 1989, Bronson 2009). Long-term field studies have found that while photoperiod controls the birth season, some ungulates can delay or advance birth timing by weeks according to precipitation and temperature variability for year-specific optimal conditions (<u>Renaud et al. 2019</u>, <u>Moyes et al. 2011</u>. <u>Berger 1992</u>, Friebe et al. 2014). In contrast, other animals heavily depend on photoperiod to time reproduction (e.g. roe deer, *Capreolus capreolus*), leaving them with little phenotypic plasticity to adjust birth timing to changing environmental conditions (<u>Rehnus et al. 2020</u>). When photoperiod is stable near the equator, timing of birth in some African animals is associated with annual patterns of rainfall and temperature, while others show no seasonality in births (<u>Ogutu et al. 2015</u>, <u>Abrahms et al. 2022</u>).

Comparative quasi-experimental studies using captive birth data offer a valuable approach to study the environmental and physiological cues of reproductive timing (<u>Clauss et al. 2021</u>, <u>Zerbe et al. 2012</u>). Most studies have been on captive ungulate species, and few studies have been focused on captive carnivore species. Some ungulate species that are seasonal in the wild become aseasonal in captivity when given year-round food, suggesting reproduction may be tied to body condition (<u>Skinner et al. 2002</u>, <u>Zerbe et al. 2012</u>). Additionally, translocating animals to zoos at different latitudes can test the role of photoperiod on reproductive timing. Some animals from the Northern Hemisphere shift their reproductive timing by six months when moved to the Southern Hemisphere, suggesting photoperiod may play a key role in reproductive timing (Marshall 1937, <u>Duke of Bedford and Marshall 1941</u>). In contrast, some tropical species maintain their original breeding season even when moved to higher latitudes, suggesting they rely on cues other than photoperiod (<u>Duke of Bedford and Marshall 1941, Porton et al. 1987</u>).

Gray wolves are among the most widely distributed land mammals, ranging from 12°N to 83°N latitude across diverse habitats. This makes them an ideal species to study how environmental variation influences timing of birth over a large spatial scale. Wolves are monoestrus and generally breed in late winter and give birth during the spring (Nagashima and Songsasen 2021, <u>Mech 2002</u>). An exception are wolves from India that breed in autumn and give birth in the winter (<u>Jhala 2003</u>). Reproduction in northern latitudes coincides shortly before the reproduction of their prey, allowing access to neonate calves during pup raising season. However, whether this is the adaptive explanation of reproductive timing remains unclear, as in some regions the highest biomass acquisition is in winter months, rather than during pup season (Gable et al. 2024). Similarly unclear is the proximate mechanisms that cue reproductive timing in wolves. While reproductive timing is likely partially controlled by photoperiod, there have been no experimental study to directly test this (Nagashima and Songsasen 2021). Removing the pineal gland in ten wolves and oral melatonin supplementation did not impact reproductive timing, however it is unclear whether there are other contributing factors (Asa et al. 1987, Kreeger et al. 1991). In captivity, a wolf pack from Indiana at 40.5°N retained the same seasonal birth timing as wild wolves at the same latitude, suggesting wild and captive wolves may be using similar environmental cues for reproduction (<u>Lord et al. 2013</u>). Overall, both the adaptive significance and physiological mechanisms of wolf reproductive phenology remains poorly understood.

Here, we investigate the environmental factors associated with reproductive phenology in a widespread large carnivore, the gray wolf (*Canis lupus*). Using 1,261 estimated birth dates from wild and captive wolves, we address three questions: (1) how does timing of birth change across latitude in wild wolves? (2) what environmental variables (daylength, temperature, precipitation, NDVI) best explain timing of birth in wild wolves? (3) how does timing of birth in captive wolves compare to wild wolves at the same latitudes?

## Methods

### Reproductive data

We compiled a dataset of estimated birthing dates from published and unpublished sources for wild wolf individuals across North America and Eurasia (Table S1). For wild wolves, we estimated the timing of birth from different sources of information based on the (1) movement patterns of the female wolf using satellite-collared and radio-collared individuals and (2) pup age based on morphological and developmental milestones through camera trap photos, observation, and pup handling. Table S2 has a full description of data types and criteria used to estimate timing of birth (Table S2).

For captive wolf data, we accessed the Zoological Information Management System (ZIMS) and recorded the birth timing and latitude at which the wolf individual gave birth. We only used data from zoos in Europe and Asia described as *Canis lupus lupus*, *C. l. signatus, C. l. arabs*, and *C. l. pallipes*. We cross-checked the *C. l. pallipes* entries with the National Studbook of the Indian wolf to verify which individuals originated in India, and therefore belong to the Indian wolf lineage, which is the earliest diverged wolf lineage (Wildlife Institute of India, 2018, Hennelly et al. 2021). We also included birth dates provided in the National Pedigree Book of the Tibetan wolf (Wildlife Institute of India, 2009). Table S3 provides a full description of the populations and subspecies used in our captive wolf dataset (Table S3). Lastly, we conducted a literature search on Google Scholar to record the season or birth timing for wild wolves. From our literature search, we recorded the date range of birth for wolf populations in Armenia, Azerbaijan, Turkmenistan, Ukraine, Belarus, and Russia (Geptner 2001, Kirilyuk et al. 2021, Sidorovich and Rotenko 2019, Table S4).

We transformed the birth timing to an adjusted Date of Year (DOY) where we set November 1^st^ as day 1 to account for the early birth times in Indian wolves. We estimated the mating date for each wolf individual’s birthing time by subtracting the average gestation length of gray wolves (65 days; <u>Concannon et al. 1983</u>, <u>Seal et al. 1979</u>) by the timing of birth.

### Environmental data

We collected nine environmental variables from Worldclim v2 at at 1×1km resolution for each wild wolf individual location: annual mean temperature (BIO1), mean diurnal range (BIO2), temperature annual range (BIO7), mean temperature of wettest quarter (BIO8), mean temperature of driest quarter (BIO9), mean temperature of warmest quarter (BIO10), mean temperature of coldest quarter (BIO11), annual precipitation (BIO12), and precipitation seasonality (BIO15) ((<u>Fick and Hijmans 2017</u>). We also calculated mean and standard deviation of the Normalized Difference Vegetation Index (NDVI) from MODIS Terra Moderate Resolution Imaging Spectroradiometer Vegetation Indices data for 2022, and filtered for clouds using R package MODIStools (Hufkens 2023). We calculated daylength at the estimated mating date using the daylength function in the package “geosphere” (Hijmans 2024).

### Statistical analysis

We used one-way ANOVA and Tukey Honest Significant Differences (HSD) tests to assess whether mean birth timing differed between our six wild wolf populations (Indian, Iberian, Northern Europe, Tibetan, North America, Southwest Asia). We grouped wolves within these populations as the following: India (India), Iberian (Spain, Portugal), Northern Europe (Sweden, Norway, Poland), Tibetan (India), North America (Canada, USA), Southwest Asia (Iran, Saudi Arabia). We tested linear, asymptotic, and logistic models using the nls2 package in the statistical program R (Grothendieck 2024, R Core Team, 2020) to assess the relationship of birth timing to latitude, with calculating marginal R^2^ using the package MuMIn in R (Barton 2024).

We used generalized linear mixed-effects models (GLMM) to evaluate which environmental factors explain timing of birth in wild wolves using the package lme4 (Bates et al. 2015). We identified correlated environmental variables (|r| > 0.7) using the corrplot library in R (Wei and Simko 2024), and subsequently performed principal component analysis (PCA) on eight correlated variables to reduce collinearity (Table S5, Figure S1). PC1 explained 58.4% of the cumulative variation, driven by positive values of annual mean temperature and quarter-based temperatures, and negative values of temperature annual range and standard deviation of NDVI (Figure S2). This corresponds to a positive PC1 representing warm environments with little variation in yearly temperature and NDVI, whereas a negative PC1 representing cold environments with greater variation in yearly temperature and NDVI. We used the term Temperature Index to refer to PC1 in all downstream analyses. Precipitation seasonality and mean temperature of the wettest quarter (BIO8) were correlated (<-0.7) and we chose to keep precipitation seasonality. We performed GLMM analyses to assess the relationship between birthing time and our fixed effect variables (daylength, Temperature Index, precipitation seasonality, annual precipitation, mean NDVI) using RegionID as a random effect. Fixed effect variables were standardized using “scale” in R. We performed GLMM on the whole dataset and only GPS-collared data, which have higher precision to compare to our full model that also used data from estimated pup ages. We performed GLMM using only unique individuals since the coefficient of variation per individuals was less than each population’s coefficient of variation (Figure S3). We used AIC model selection through the function “dredge” in the package MuMIn in R.

For our captive wolf dataset, we compared birth timing to wild wolves using Welch’s t-test using unequal variances. We tested linear, asymptotic, and logistic models for the relationship between birth timing and latitude. We performed a pairwise t-test within 10 degree latitudinal bins to assess whether captive and wild wolves differed in timing of birth at the same latitudinal range.

## Results

### Timing of birth in wild wolf populations

We collected 572 birth timings from 330 wild wolves from North America, Europe, and Asia. Birth timings of wild wolves ranged from November 22^nd^ for Indian wolves at 18°N to June 15^th^ for Arctic wolves at 80°N (Figure 1A). For a single population in Yellowstone National Park that we recorded 172 birth timings from 73 individuals, the standard deviation around the mean (April 18th) was +/- 8.3 days with birth timings ranging from April 1^st^ to May 7^th^. One-way ANOVA revealed significant differences in mean timing of birth among the six main wolf populations, with Turkey HSD indicating that Indian and Iberian wolves significantly differed from all other wolf populations (p-value < 0.01; Table S6). Indian wolves gave birth significantly earlier than all other wolf populations (average: January 10^th^, SD_India_= 27 days, n=43). The timing of birth in Iberian wolves occurred later than wolves at similar latitudes, with an average birth date in Iberian wolves being May 21^st^ (n=28).

**Figure 1.**
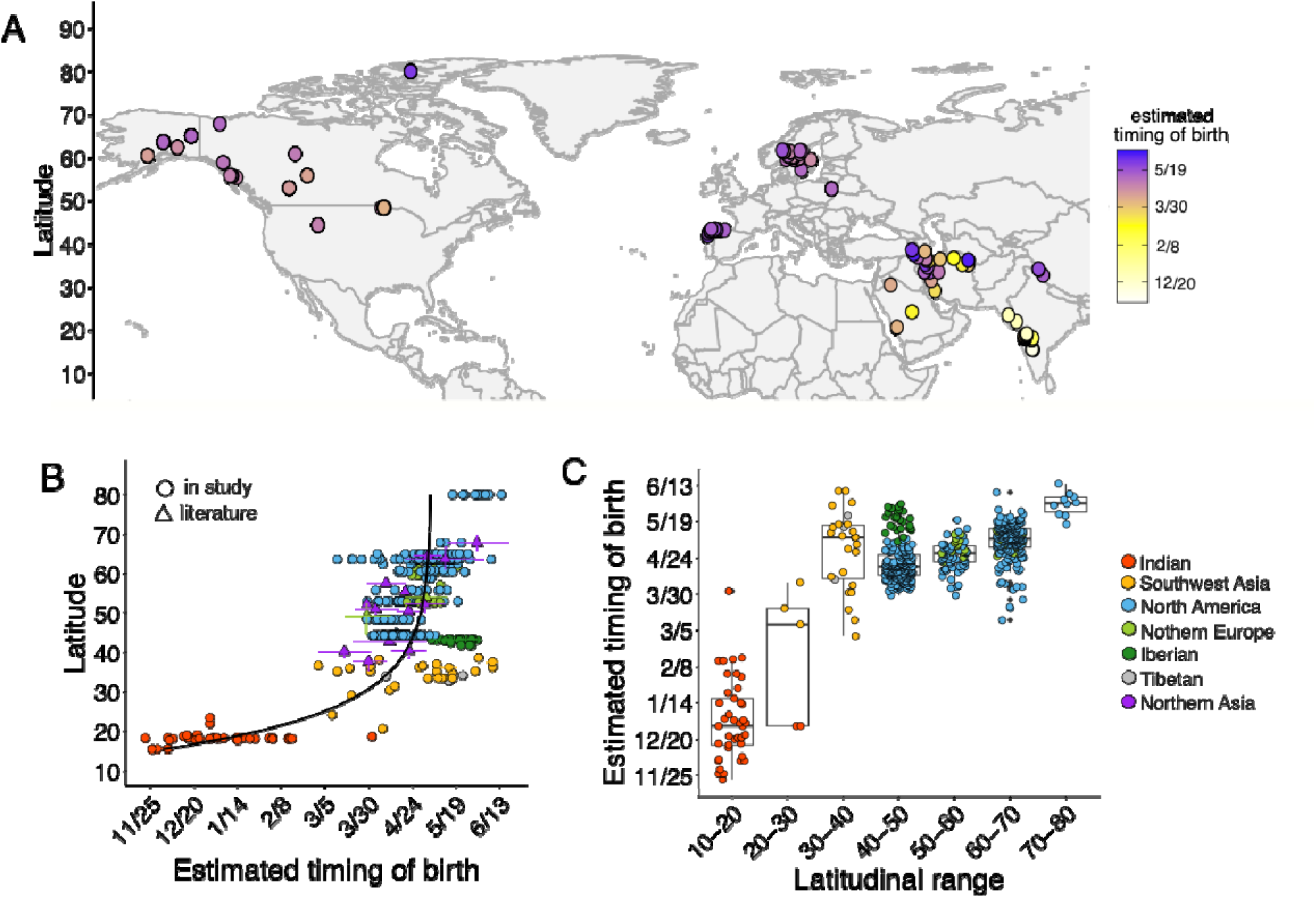
**(A)** Location and estimated timing of birth for wild wolves in Europe, North America, and Asia (n=572). **(B)** Data points for timing of birth (n=572) from 330 wild wolf individuals (circles) and documented ranges of birth timing for 16 different wolf populations (triangles) described in published literature sources (**Table S1**). The black curve indicates the asymptotic relationship between timing of birth and latitude and explains 77% of birth timing variation (pseudo R^2^=0.77). **(C)** Mean and variation of timing of birth across different latitudinal ranges for wild wolves with colors indicating population assignment. (**Table S1**).

**Figure 2.**
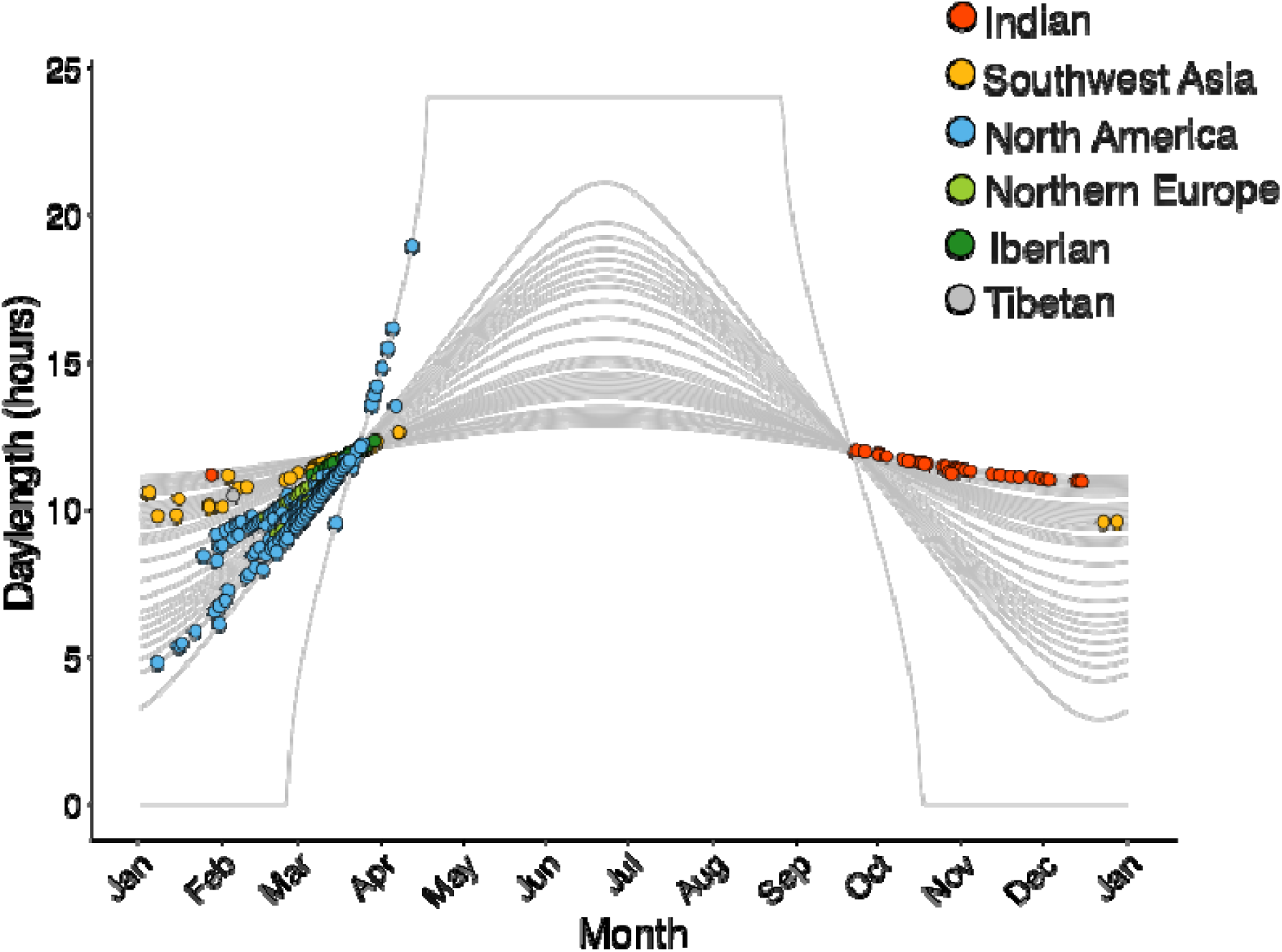
Relationship between daylength change across a single year and the estimated date of mating (n=572) for wild gray wolves at different latitudes. Circles indicate estimated timing of mating for individual wild wolves, which was estimated by subtracting the average gestation time (65 days) from the estimated timing of birth. Lines indicate the daylength change across a year for each latitude we had wild wolf data from.

Birth timing occurred later with increasing latitude following an asymptotic relationship (Figure 1C, Figure S4, Table S7). There was a greater change in birth timing with increasing latitude at lower latitudes, where the average birth timing difference was 108.5 days between 10-20N (average 63.3 DOY, n=41) and 40-50N (average 171.5 DOY, n=220) compared to an average difference of 40.3 days between 40-50N and 80-90N in northern latitudes (212 DOY, n=10). Latitude explained 77% of the birth timing variation using our asymptotic model (pseudo R^2^=0.76, RSE=17.3 days). Within wolves from Europe, birth timing had a negative linear relationship with latitude (β =-2.96, SE=3.05, df=26, p=0.34; Figure S5), driven by the late birth timing of Iberian wolves, who typically gave birth later compared to wolves in Poland, Sweden and Norway.

### Environmental factors associated with timing of birth in wild wolves

Our top GLMM model included all five fixed effect variables (daylength during mating season, precipitation seasonality, temperature index, annual precipitation, mean NDVI) (Table 1, Table S8). The next best model that excluded mean NDVI was within 2 AIC values (ΔAIC = 1.67), otherwise, none of the other 30 possible models were within 2 AIC values (Table S8). Daylength during mating season had the strongest association with birth timing (βdaylength= 9.81, SEdaylength= 0.50, Tvaluedaylength= 19.31, pdaylength=<0.0001) (Table 1), with births occurring later as daylength increased during mating season. This pattern was consistent when using only data from GPS-collars wolves (n=297) that were from Europe (n=43), and North America (n=255, Table S9), and only using unique wolf individuals (Table S10).

**Table 1.** The generalized linear mixed-effect regression model with the lowest AIC for evaluating the association between timing of birth and four environmental variables (daylength, temperature index, annual precipitation, precipitation seasonality) for 572 birth timings from 330 wild gray wolves. We included RegionID as random effects to account for population assignment of which individuals were from.

| **Fixed Effect** | **Estimate** | **SE** | **Df** | **T value** | **P value** |
| --- | --- | --- | --- | --- | --- |
| Daylength | 9.81 | 0.50 | 562.1 | 19.31 | <0.0001 |
| Temperature Index | 10.99 | 1.72 | 565.0 | 6.38 | <0.0001 |
| Annual precipitation | 3.57 | 0.68 | 563.6 | 5.23 | <0.0001 |
| Precipitation seasonality | 6.024 | 0.86 | 565.9 | 6.97 | <0.0001 |
| Mean NDVI | -1.072 | 0.83 | 564.2 | -1.28 | 0.19 |

Our GLMM model indicated precipitation-related and temperature-related variables were additional factors associated with timing of birth. Timing of birth was negatively associated with Temperature Index, in which early births were associated with warm environments with little yearly temperature and NDVI variation (Table 1). High annual precipitation variability was also associated with earlier birth timing, which was driven by the high annual precipitation variability for Indian wolves due to monsoon season. When conducting a linear regression model for each environmental variable for all wild wolves, we found daylength during mating season explained little variation in timing of birth (R^2^<0.01), whereas Temperature Index (R^2^=0.46) and Precipitation Seasonality (R^2^=0.07) explained the most (Figure S6). This pattern was entirely driven by Indian wolves; when they are removed from the analysis, daylength during mating season explained the most variation in birth timing (R^2^=0.47) (Figure S7). Unlike other wolf populations, wolves in Europe had a negative relationship between birth timing and Temperature Index, where later birth timings were associated with warmer temperatures with little yearly temperature variation (βPC1= -6.021, SEPC1= 0.82, TvaluePC1= -7.287, pPC1=<0.0001, all European wolves Figure S8). This relationship is driven by Iberian wolves, removing them resulted in no significant relationship between birth timing and Temperature Index in wolves from Europe (βPC1= 2.993, SEPC1= 2.069, TvaluePC1= 1.447, pPC1=0.161 for European wolves without Iberian wolves, Figure S12).

### Comparison between wild and captive wolves

From ZIMS, we recorded 688 birth timings from captive wolves in European and Asian zoos. Our data consisted of *Canis lupus signatus* (n=63), *Canis lupus lupus* (n=523), *Canis lupus arabs* (n=18), *Canis lupus pallipes* (n=64), and *Canis lupus chanco* (n=25). Timing of birth differed significantly between wild and captive wolves (t=6.16, df=1270, p=<0.0001), but largely fell within the range of the wild wolf dataset. Captive wolves gave birth slightly later (May 2^nd^) compared to wild wolves (April 21^st^), however, the standard deviation was much higher for the captive population (sd_captive_=40.5, sd_wild_=35.0). The higher standard deviation in captive wolves may be related to more out-of-season births, which has been documented in other canids in captivity attributed to stress and behavioral changes (Belyaev and Trut 1975). We found captive Iberian wolves gave birth later than the captive Eurasian wolves, *C. l. lupus* when at the same latitude (Figure S13). For example, at the latitudinal range of the Iberian peninsula (33-43°N), the average timing of birth for captive Iberian wolves was June 5^th^ (n=40), compared to May 24^th^ for captive *C. l. lupus* at this latitudinal range (n=17). At the equator around 1-10°N, wolves gave birth across multiple months (Figure 3A).

**Figure 3.**
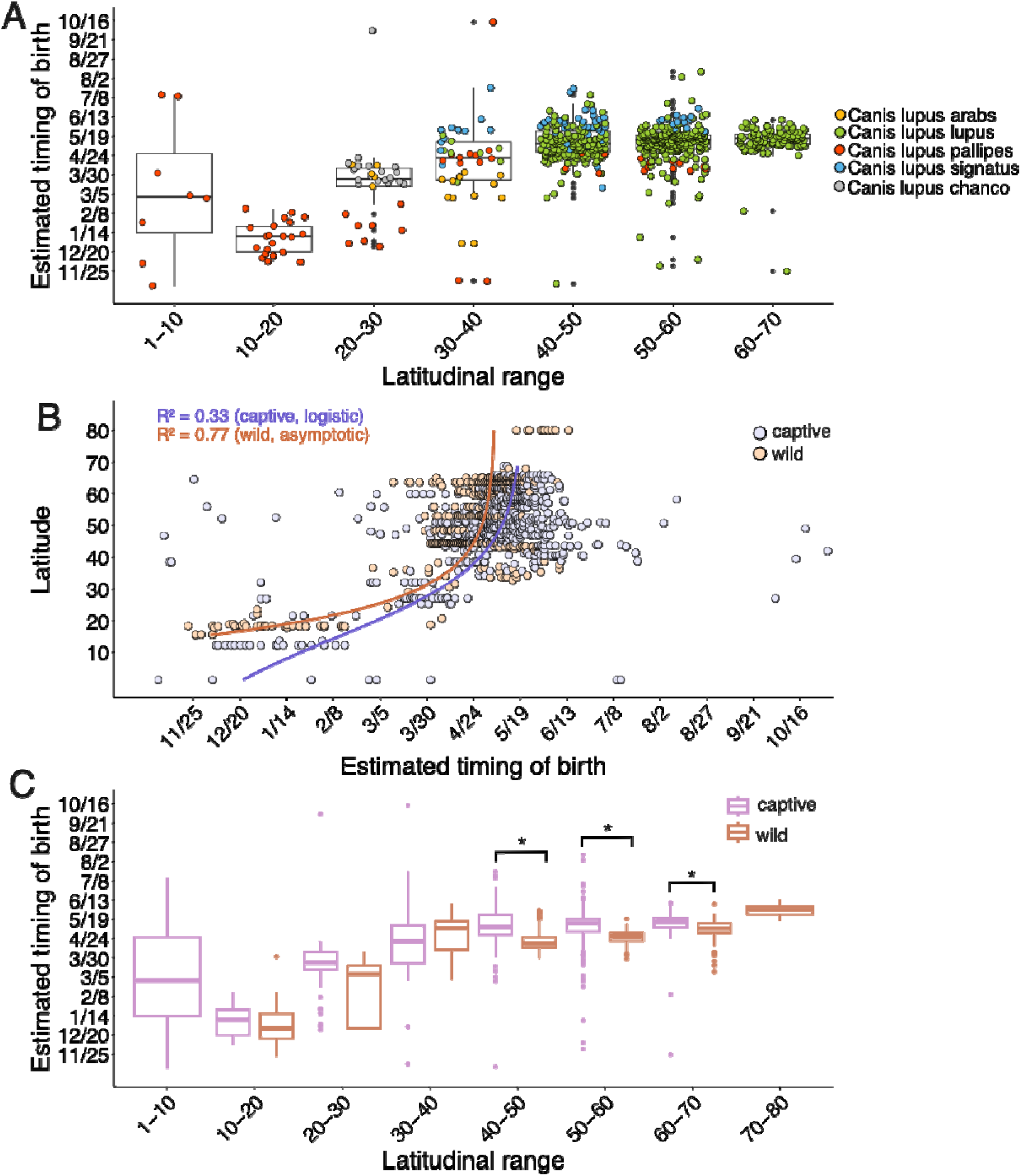
**(A)** Mean and variation of timing of birth within different latitudinal ranges for 668 captive wolf individuals. Colors indicate the assigned subspecies provided within the ZIMS database **(B)** Relationship between latitude and estimated timing of birth for captive wolves (orange) and wild wolves (purple). For captive wolves, latitude explains 33% of the variation in the timing of birth using a logistic model, whereas for wild wolves, latitude explains 77% of the variation in timing of birth using an asymptotic model. **(C)** Comparison for the mean and variation of timing of birth for 668 captive wolves and 572 wild wolves. Only wolf populations in the latitudinal range between 40-50°N, 50-60°N and 60-70°N were statistically different between the timing of birth of captive and wild wolves.

When considering how variation in birth timing relates to latitude, birth timing in captive wolves was similar to wild wolves. For captive wolves, birth timing occurred later with increasing latitude following a logistic relationship (Table S7, Figure S14). The logistic model explained 33% of the variation in timing of birth (pseudo R^2^=0.33), which was less than for wild wolves (pseudo R^2^=0. 77) (Figure 3B). We found the variation of birth timing within each ten degrees of latitude is similar between captive and wild wolves (Figure 3C), with only the timing of birth at 40-50°N, 50-60°N and 60-70°N significantly differ (p<0.05) between captive and wild wolves.

## Discussion

Reproductive phenology is a fundamental aspect of an organism’s life history with direct impacts on reproductive success. Understanding the environmental cues that organisms use to time reproduction is therefore crucial for predicting species’ responses to climate change. Wild gray wolves have a seven-month difference in reproductive phenology, from birth occurring in mid-November at 18°N for Indian wolves to mid-June at 80°N for Arctic wolves. We found photoperiod is strongly associated with reproductive phenology for wolves. Climatic-related variables also contribute to explaining reproductive phenology, especially in lower latitudes when photoperiodic cues are more stable. This suggests populations across their range may rely on different environmental cues to time reproduction, which has important implications for predicting how climate change may differentially impact populations. Our work adds to the limited number of studies on large carnivore reproductive phenology and improves predictions of climate-induced shifts for higher trophic levels (<u>Garcia-Rodriguez et al. 2020</u>, <u>Mahoney et al. 2020</u>, <u>Mattisson et al. 2022</u>, <u>Abrahms et al. 2022</u>).

Multiple lines of evidence suggest reproduction in gray wolves is partially physiologically linked with photoperiod, similar to other temperate canids like red fox (*Vulpes vulpes*) and arctic fox (*Vulpes lagopus*) (Kuznetsov 1979, <u>Forsberg et al. 1989</u>, <u>Forsberg et al. 1990</u>). First, captive wolves in Tsitsikamma, South Africa shift reproduction by six months, giving birth to pups in the autumn months (Tsitsikamma Wolf Sanctuary, pers. comm.). Tsitsikamma is at 33°S with a mild climate and little yearly temperature and precipitation variation, suggesting increasing daylength during autumn may be the primary cue for reproduction. Second, our data suggests captive Indian wolves are aseasonal when translocated to 1.5°N in Singapore, where there is no daylength change and little annual variability in temperature and rainfall (Figure 3A, <u>Wildlife Institute of India 2018</u>). Lastly, captive wolves assigned as *Canis lupus pallipes* that presumably originated from 13°N-41°N delayed their reproductive season when translocated to a higher latitude (Figure 3A).

The adaptive significance of wolf reproductive phenology could be related to the energetic demands of acquiring sufficient food for pups with limited mobility. In particular, wolves generally give birth shortly before birth pulses of their ungulate prey, which occurs around peak forage productivity in the spring (<u>Stoner et al. 2016</u>, Post 2003, Michel et al. 2020). We observe this for Indian wolves giving birth before their prey blackbuck (May, <u>Priyadarshini et al. 2022</u>), wolves in Yellowstone with elk (late May, Barber-Meyer et al. 2008), wolves in Sweden and moose (late May, <u>Neumann et al. 2020</u>) Arctic wolves with Peary caribou (early June in Western Greenland, <u>Post and Forchhammer 2007</u>). Birth pulses of ungulates are a predictable resource each year, with the smaller bodied neonates being easily captured and transported to den sites.

However, the pup-rearing season, when the energetic demands of adult wolves are highest, is not when biomass acquisition of wolves is the highest (Gable et al. 2024, Metz et al. 2012). Instead, biomass acquisition typically peaks in winter when ungulates are in poorest body condition (Nicholson et al. 2008, Desforges et al. 2020). In some systems, biomass acquisition of adult wolves often reaches its minima of the annual cycle during or shortly after ungulate birth pulses (Metz et al. 2012, Voyageurs Wolf Project unpublished data) while energetic expenditure likely reaches its maxima, with wolves expending up to 17-20% more energy per day during the pup-rearing season than other times of the year (Bryce et al. 2022). Consequently, starvation during summer in many systems is one of the most common causes of mortality for wolf pups (Benson et al. 2013, Hynes et al. 2026, Gable et al. 2023).

Although reproductive phenology appears well-timed to exploit ungulate birth pulses, there may be other adaptive explanations. In northern latitudes, climatic conditions may play an important role for reproductive success in rearing young pups and for the lactating female wolf. While wolf pups can withstand inclement weather and cold to a certain point, having pups during extended periods of such conditions may exceed their physiological tolerance. In addition, end of gestation and early lactation are often the most nutritionally demanding for mammals (<u>Clutton-Brock et al. 1989</u>). For dogs, end of gestation can require 25-50% above maintenance energy requirements and lactation can require up to 2.5-3 times more maintenance energy during the first postpartum month (<u>Fontaine 2012</u>). Lactating wolves may increase their water and food intake by two to three times more compared to a non-lactating wolf (Oftedal and Gittleman 1989). Correspondingly, wolf dens are often located nearby fresh water sources (<u>Trapp et al. 2008</u>, <u>Iqbal et al. 2025</u>, <u>Khan et al. 2024</u>, <u>Habib and Kumar 2007</u>). For northern latitudes, peak biomass accumulation in the winter months coincides with late gestation for the female wolf, which may suggest energy intake during this period could be more important than during pup rearing. Lastly, a reliance on photoperiod and the gestation window (∼65 days) may physiologically constrain birth timing to a specific seasonal window independent of what might be theoretically better suited. Further work with empirical data to test these predictions would help understand the tradeoffs of wolf reproductive phenology and its evolution in grey wolves.

Our wide latitudinal sampling allowed us to explore how wolves have evolved to time reproduction when photoperiodic cues are more stable near the equator. Despite little annual daylength change at 10-20°N, we found reproductive phenology remained seasonal. For instance, wild wolves from central India retained birth seasonality with most births occurring in December and January. While our data for Indian wolves was primarily from camera trap photos and may not be as accurate as GPS-collared data, our results align with previous reported breeding times based on observations and monitoring over multiple years (Jhala 2003, M.G., S.K., B.H., Y.V.J., pers. comm.). Captive Indian wolves showed the same seasonal timing, suggesting body condition is not a primary factor. We suspect Indian wolves are either responding to very subtle daylength changes during the mating season or rely on climatic cues, such as related to temperature or precipitation. Seasonal reproduction in areas where photoperiodic cues are stable has also documented in other canid species, such as Ethiopian wolves *Canis simensis* (10°N; late August-December), black-backed jackals *Canis mesomelas* (2.5°S, July-September), and African wolves *Canis lupaster* (2.5°S, December-March) (Moehlmann 1983, <u>Moelhman 1986</u>, Silero-Zubiri and Gottelli 1994, Sillero-Zubiri et al. 1998). For these canids, the presence of a breeding season suggests that social or climatic cues, such as changes in rainfall or temperature, may trigger reproduction. We hypothesize Indian wolves may use climatic cues, particularly the mating season may be associated by reduced rainfall or cooler temperatures as the end of the monsoon season in early October. Recent work has also identified secondary contact zones with little gene flow between Indian wolves and other nearby wolf lineages (Hennelly et al. 2024, Hennelly et al. 2026). We hypothesize difference in reproductive phenology may contribute to reduced gene flow between the Indian wolves and nearby Tibetan wolves and in western regions of Pakistan and Iran. These findings support broader work suggesting locally adapted reproductive phenologies may be more prevalent in tropical regions and play an important role in population divergence (Moore et al. 2005).

Interestingly, we found Iberian wolves consistently reproduce later than expected based on latitude. Our data on Iberian wolves comes from three separate studies that show similar results, two of which estimated birth timing from GPS-collared wolves and one from young pups that were removed from dens (Figure S15). This pattern was observed in captivity, where captive Iberian wolves translocated to non-native (>44°N) higher latitudes showed delayed reproduction compared to captive Eurasian wolves (*Canis lupus lupus*) (Figure S13). We hypothesize delayed birth timing in Iberian wolves could be associated with climatic differences of the Iberian peninsula. Unlike continental Europe and Central Asia, the northern Iberian peninsula has a mild climate with consistent rainfall that peaks in winter, and does not experience freezing (<0°C) or hot temperatures (>26°C). In northern Iberian peninsula, calving season for ungulates falls between April and May, aligning with late gestation and pup rearing season (<u>Lopez et al. 2016</u>). We also observe high variability in birth timing in Iranian wolves during spring months. This could be due to a more stable photoperiod, differences in climatic factors compared to northern latitudes, due to our data being from camera trap photos. Confirming with GPS-collared individuals would be needed to assess whether this variability is reflective of reproductive phenology of Iranian wolves. Population-specific reproductive timing has been documented in other mammals and birds using common garden or genetic studies (<u>Rosmalen et al. 2023</u>, <u>Fudicker et al. 2016</u>, <u>Gwinner and Scheuerlein et al. 1999</u>, <u>Dark et al. 1983</u>). Future work to understand how much reproductive timing is genetically vs. environmentally driven has broad implications for predicting evolutionary responses to climate change (<u>Bradshaw and Holzapfel 2006</u>).

Our work on wolf reproductive phenology has important conservation and management implications. As climate change alters the timing of resource availability in many places, animals must adjust their reproductive phenology or else risk a phenological mismatch (Visser and Gienapp 2019). In North America, previous work found wolf den timing did not shift over a 17 year period, whereas annual start of the growing season shifted 8 days earlier (Mahoney et al. 2020). This pattern could be consistent with wolves in northern latitudes primarily relying on photoperiod, which may restrict their ability to fine-tune reproductive timing according to yearly shifts in annual temperature and rainfall. We highlight another potential phenological mismatch for Indian wolves that may make them vulnerable to the effects of climate change. Indian wolves give birth just before the hottest and driest time of the year (March to May). Heat waves during these months are expected to increase and become earlier due to climate change, with temperatures reaching as high as 50°C (122°F) for multiple days (<u>Rohini et al. 2019</u>). During this time, the lactating female wolf may be at significant risk of dehydration and high temperatures may reduce the packs’ ability to forage. For African wild dogs, high temperature during denning is associated with reduced foraging times and reproductive success (<u>Woodroffe et al. 2017</u>). Second, if Indian wolves use climatic cues to time reproduction, the increasing unpredictability of the monsoon season could cause maladaptive shifts in birth timing (<u>Sandeep et al. 2018</u>). A phenological mismatch has been documented in African wild dogs where climate-driven shifts in breeding timing increased exposure to high temperatures during the denning season and reduced reproductive success (<u>Abrahms et al. 2022</u>). Establishing long-term monitoring programs for Indian wolves may be important to assess how a changing climate may impact reproductive success and population persistence. Lastly, we highlight that wolf-dog hybridization may have important implications on wolf reproductive phenology. Dogs are no longer seasonal reproducers and, if aseasonality is inherited, could alter breeding times in admixed wolves, a possibility that has been suggested for wolves in Belarus (<u>Sodorovich and Rotenko 2019</u>).

Finally, understanding reproductive phenology at higher trophic levels has far-reaching implications for predicting trophic-wide responses to climate change and ecosystem dynamics. Whether large carnivores are able to adjust reproductive timing to follow changing optimal conditions due to climate change will have consequences on prey communities and the broader ecosystem (Thackeray et al. 2016). The gray wolf’s distribution and evolutionary history occurred mostly in northern latitudes where photoperiodic cues are most reliable for aligning reproduction with primary productivity (<u>Sotnikova and Rook et al. 2010</u>). Similarly, many large ungulates in northern latitudes have to evolved to use photoperiodic cues for reproductive phenology (<u>Lu et al. 2010</u>, Sempere et al. 1993, Verme et al. 1987, Rolf and Fischer et al. 1996). This pattern, in which animals that primarily evolved at higher latitudes have reproduction partially cued by photoperiod, is described for ungulates and other carnivores, and may be true for some canids (<u>Duke of Bedford and Marshall 1941</u>, <u>Heldstab et al. 2018</u>, <u>Bronson 1988</u>, <u>Porton et al. 1987</u>). In such cases, predator and ungulate prey communities may have evolved reliance on similar environmental cues, such as photoperiod, allowing for predators to reliably time reproduction with their prey. However, species may differ in how they respond to shifting environmental cues under climate change and in the plasticity of those responses, potentially disrupting time-based species interactions across trophic levels (Miller-Rushing et al. 2010, Yang and Rudolf 2009). Whether trophic communities synchronized with photoperiod in temperate regions or ones that rely on climate-driven cues in tropical regions are more vulnerable to such disruptions remains an important question. Overall, our work suggests grey wolves use multiple environmental cues to time reproduction in different parts of their wide distribution. The use of different environmental cues has important implications for considering population-specific responses to climate change and may have facilitated the gray wolf’s adaptation to diverse environments and latitudinal ranges.

## Acknowledgements

L.M.H. thanks the National Science Foundation Postdoctoral Research Fellowship (award number 2208950) for funding and support. JVLB thank the Spanish Ministry of Science and Innovation (PID2023-149634OB-I00), the Spanish Ministry for the Ecological Transition and the Demographic Challenge and the Regional Government of Asturias. He also thanks to Emilio J. García for his effort in fieldwork activities. We thank Iravatee Majgaonkar for efforts with the Indian wolf breeding data. We thank Laurie Bingaman Lackey and Andrew Kitchener for assistance with accessing the captive wolf dataset.

## Data Availability Statement

Raw data and analysis output is archived in a Dryad digital repository available at https://doi.org/10.5061/dryad.sqv9s4nj5. Raw data, analysis output, and code is provided at https://github.com/hennelly/Wolf_Reproductive_Phenology

## Author contributions

**LMH** conceived study, curated the data, performed final statistical analyses, data visualization, and wrote the first draft of the manuscript. **SS** performed data collection, curated data, and performed exploratory data analysis. **BH** performed data collection and curated data, **SK, AM, MF, LL, JB, AH, AW, MZI, YVJ, JVLB, GR, MG, IG, HS, CW, PW, BZ, PM, DS, ES, MM, KC, TG** performed data collection, all authors contributed to review and editing of the manuscript.

