## Supplementary material for "Population-specific drivers of reproductive phenology in a widespread large carnivore, the gray wolf": Figure S1

**Supplemental Figures**

**Figure S1**. Pearson’s correlation coefficient between all 12 variables in the study, in which eight variables had a correlation coefficient above 0.7 (**Table S5**).

**Figure S2**. Principle component analysis of eight environmental variables.

**Figure S3**. Coefficient of variation for each individual that we have multiple recorded timing of birth for multiple years (circles) compared to the coefficient of variation within each of their respective population (bars).

**Figure S4**. Comparison between three models (linear, asymptotic, and logistic models) for assessing the relationship between adjusted DOY and latitude for wild wolves (n=572 timings).

**Figure S5.** Relationship between estimated timing of birth and latitude in which the female wolf gave birth

**Figure S6**. The relationship between estimated timing of birth for wild wolves (n=572) and each environmental variable in our dataset, including latitude

**Figure S7**. The relationship between estimated timing of birth for wild wolves without Indian wolves and each environmental variable in our dataset, including latitude

**Figure S8**. The relationship between estimated timing of birth for wolves from Europe and Iberian peninsula each environmental variable in our dataset, including latitude

**Figure S9**. The relationship between estimated timing of birth for wolves from Southwest Asia (Iran, Saudi Arabia) and each environmental variable in our dataset, including latitude

**Figure S10**. The relationship between estimated timing of birth for Indian wolves and each environmental variable in our dataset, including latitude

**Figure S11**. The relationship between estimated timing of birth for North American wolves and each environmental variable in our dataset, including latitude

**Figure S12**. The relationship between estimated timing of birth for European wolves without wolves from Iberian peninsula and each environmental variable in our dataset, including latitude

**Figure S13.** Timing of birth between 35-55N for captive wolves assigned as Eurasian wolves (*Canis lupus lupus*) and Iberian wolves (*Canis lupus signatus*).

**Figure S14**. Comparison between three models (linear, asymptotic, and logistic models) for assessing the relationship between adjusted DOY and latitude for captive wolves (n=688 timings).

**Figure S15.** Timing of birth of our data on wild Iberian wolves (*Canis lupus signatus*). This consists of 29 timings of birth that were estimated using GPS-collared wolves and estimating age of wolf pups that were removed from dens when pups were less than 15-20 days old

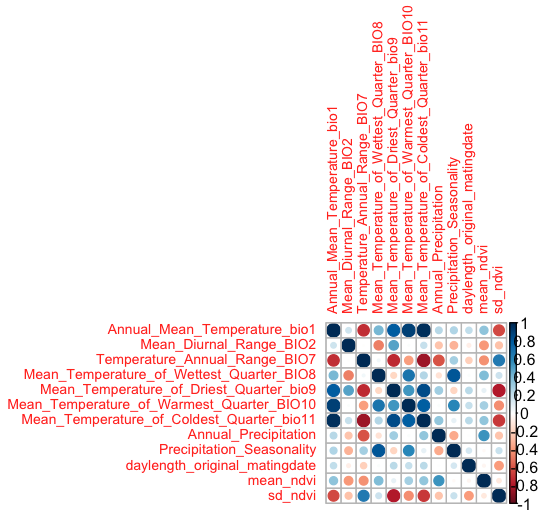

**Figure S1**. Pearson’s correlation coefficient between all 12 variables in the study, in which eight variables had a correlation coefficient above 0.7 (**Table S5**).

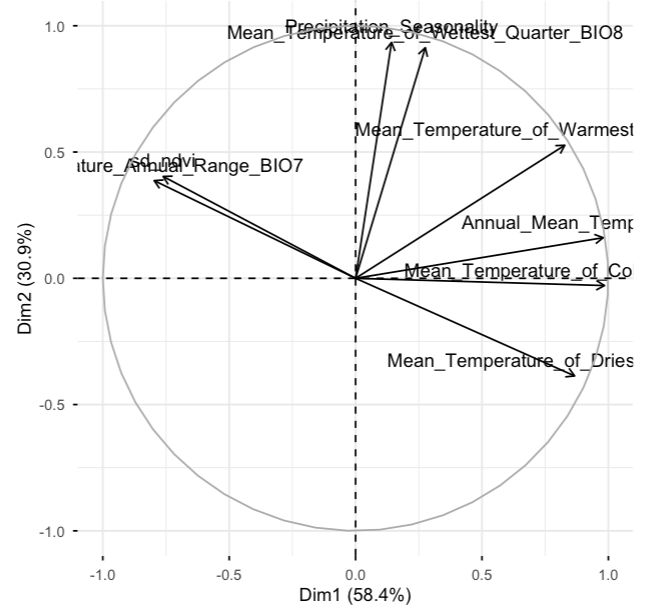

**Figure S2**. Principle component analysis of eight environmental variables, in which PC1 is largely driven by positive values of annual mean temperature and quarter-based temperatures, and negative values of the temperature annual range and the standard deviation of NDVI.

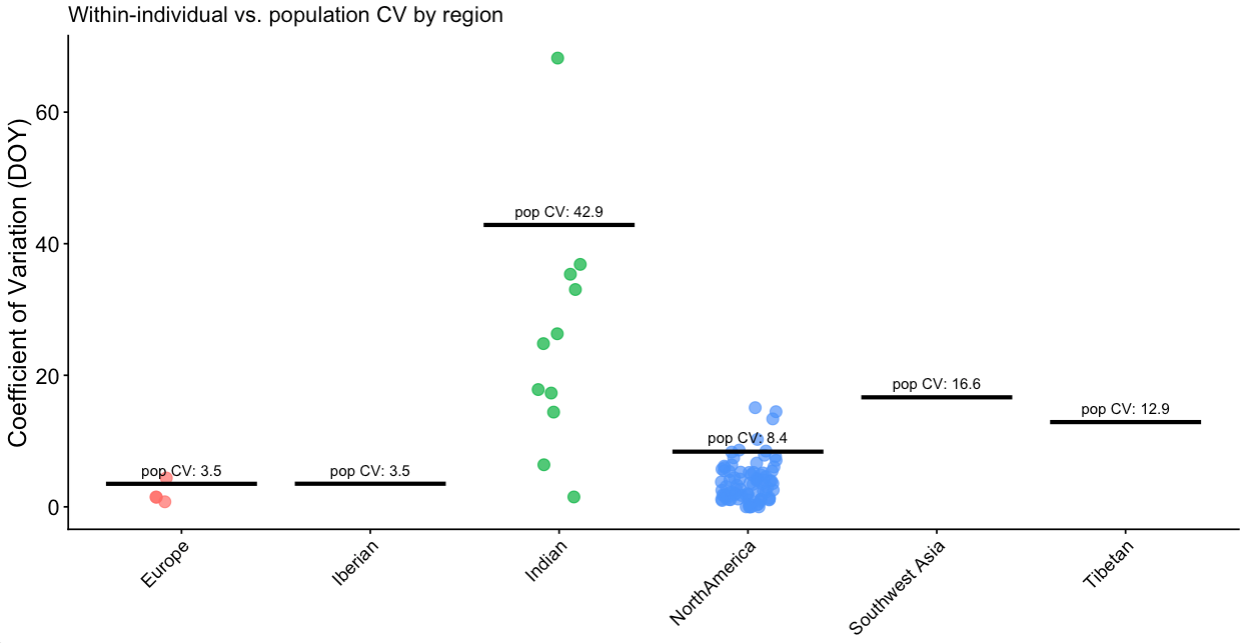

**Figure S3**. Coefficient of variation for each individual that we have multiple recorded timing of birth for multiple years (circles) compared to the coefficient of variation within each of their respective population (bars).

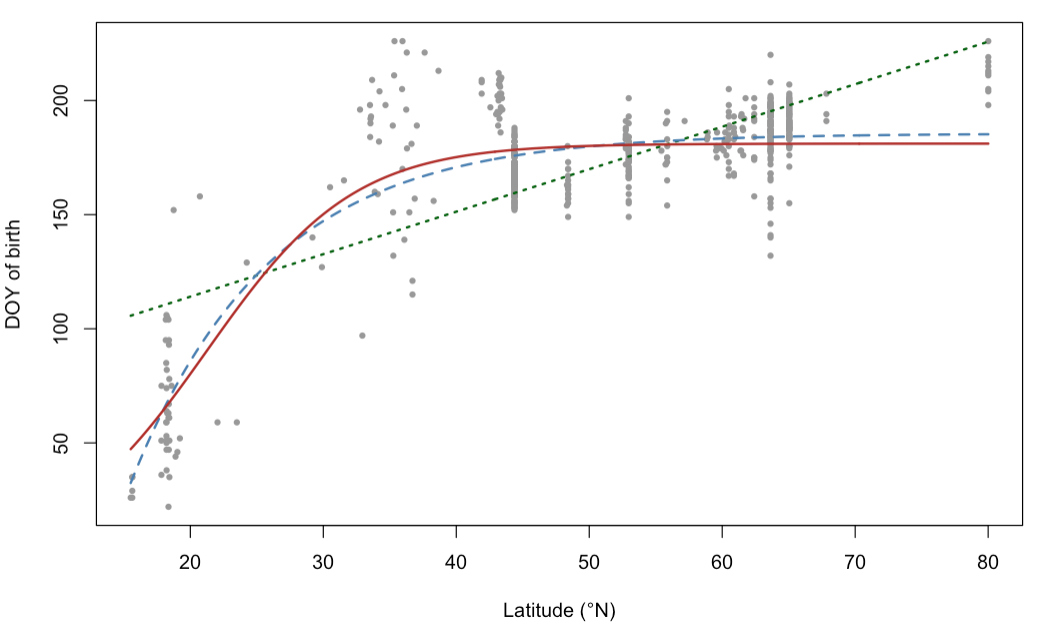

**Figure S4**. Comparison between three models – linear (green), asymptotic (blue), and logistic (red) – for assessing the relationship between adjusted DOY and latitude for wild wolves (n=572 timings). The results of the three models are found in **Table S7**.

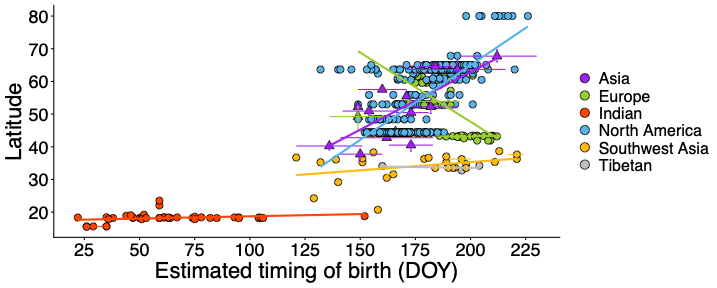

**Figure S5.** Relationship between estimated timing of birth and latitude in which the female wolf gave birth. Datapoints for timing of birth (n=572) for 330 wolf individuals (circles) and documented ranges of birth timing for 16 different wolf populations (triangles) described in published literature sources. The linear regression line is shown for each geographic region. DOY is adjusted where November 1^st^ is Day 1.

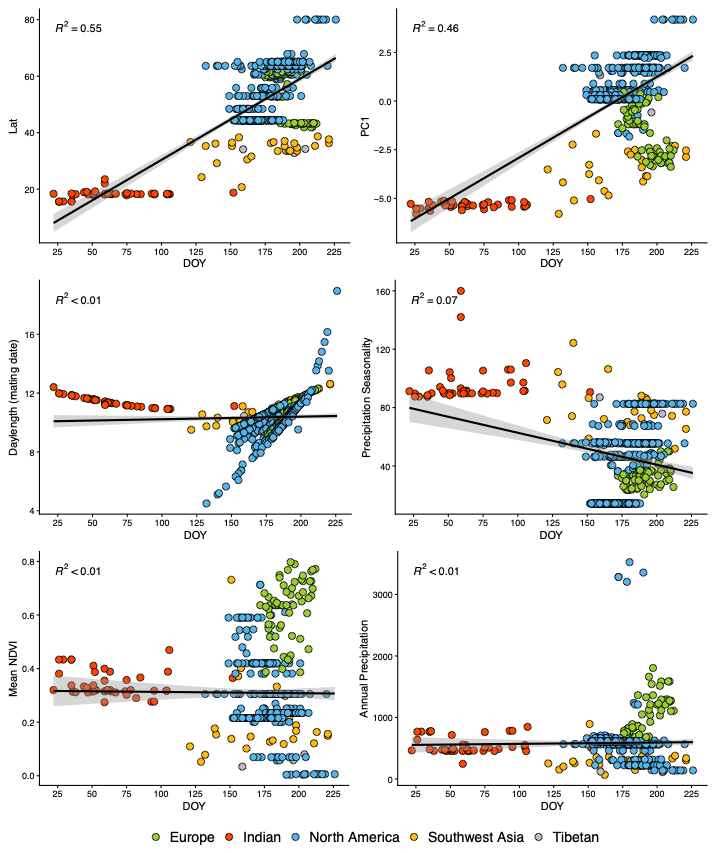

**Figure S6**. The relationship between estimated timing of birth for wild wolves (n=572) and each environmental variable in our dataset, including latitude. The linear regression line is shown for each relationship and the R^2^ value. DOY is adjusted where November 1^st^ is Day 1.

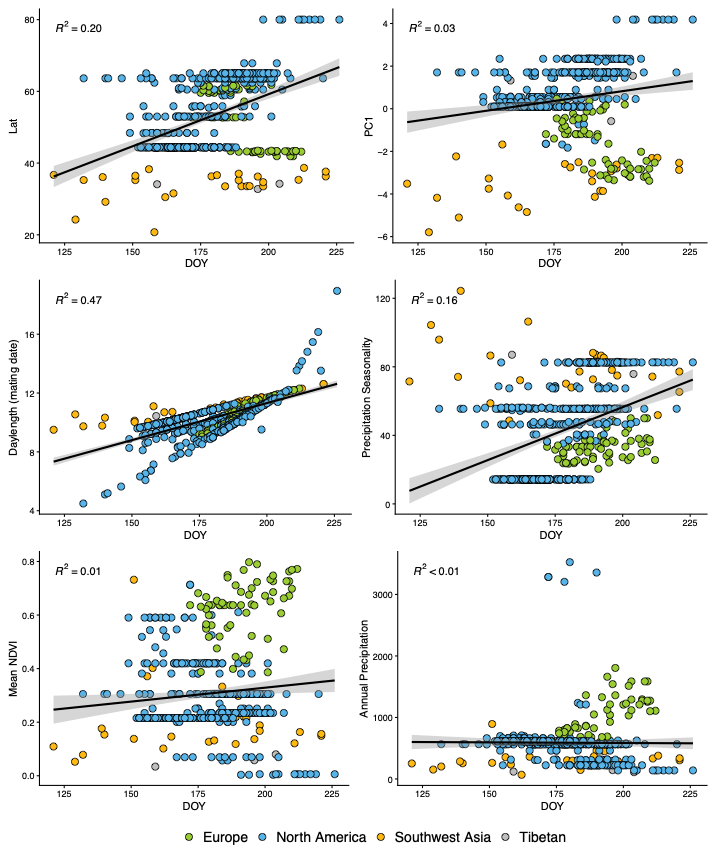

**Figure S7**. The relationship between estimated timing of birth for wild wolves without Indian wolves and each environmental variable in our dataset, including latitude. The linear regression line is shown for each relationship and the R^2^ value. DOY is adjusted where November 1^st^ is Day 1.

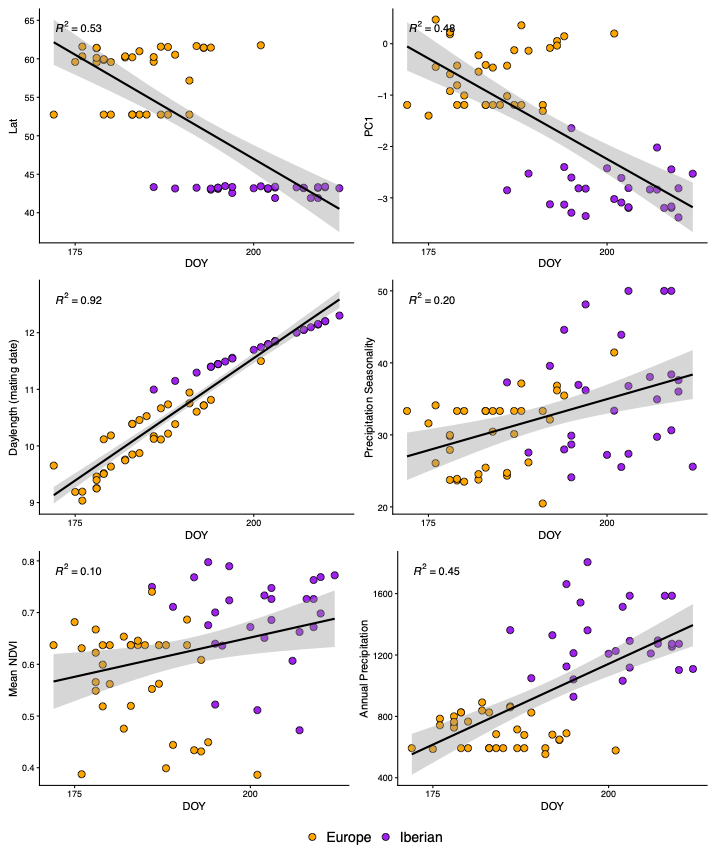

**Figure S8**. The relationship between estimated timing of birth for wolves from Europe and Iberian peninsula each environmental variable in our dataset, including latitude. The linear regression line is shown for each relationship and the R^2^ value. DOY is adjusted where November 1^st^ is Day 1.

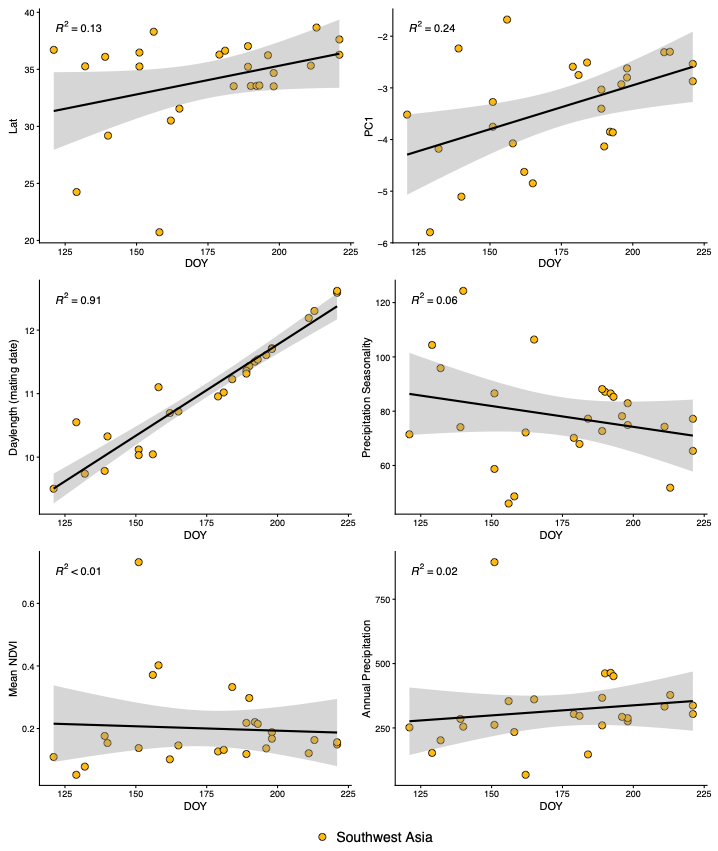

**Figure S9**. The relationship between estimated timing of birth for wolves from Southwest Asia (Iran, Saudi Arabia) and each environmental variable in our dataset, including latitude. The linear regression line is shown for each relationship and the R^2^ value. DOY is adjusted where November 1^st^ is Day 1.

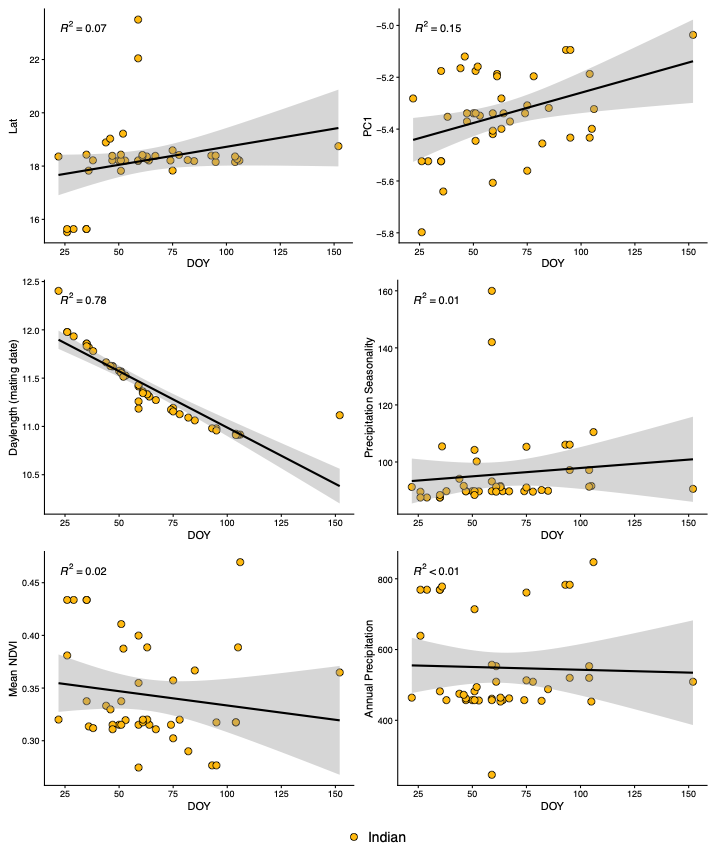

**Figure S10**. The relationship between estimated timing of birth for Indian wolves and each environmental variable in our dataset, including latitude. The linear regression line is shown for each relationship and the R^2^ value. DOY is adjusted where November 1^st^ is Day 1.

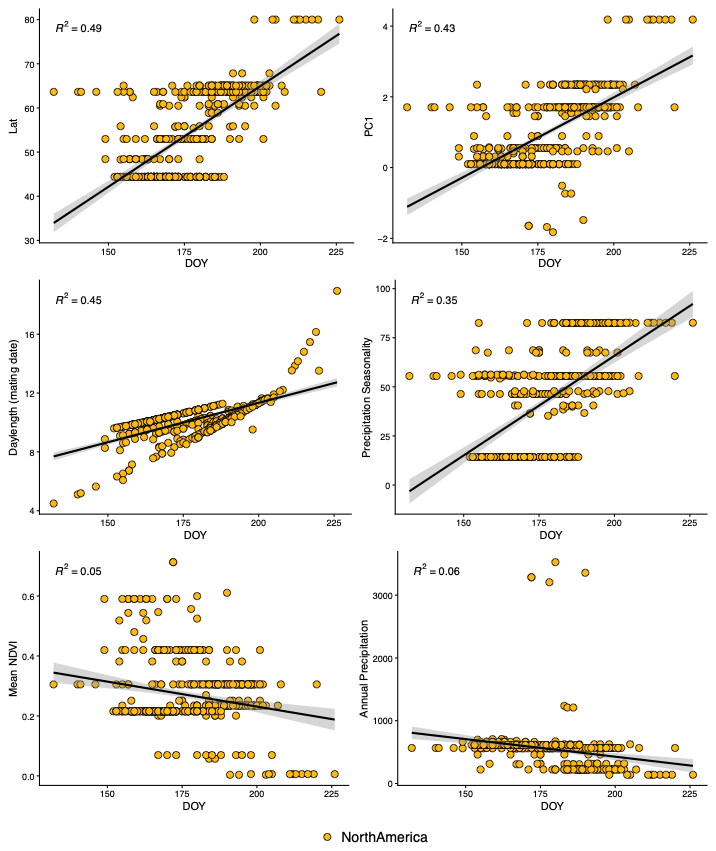

**Figure S11**. The relationship between estimated timing of birth for North American wolves and each environmental variable in our dataset, including latitude. The linear regression line is shown for each relationship and the R^2^ value. DOY is adjusted where November 1^st^ is Day 1.

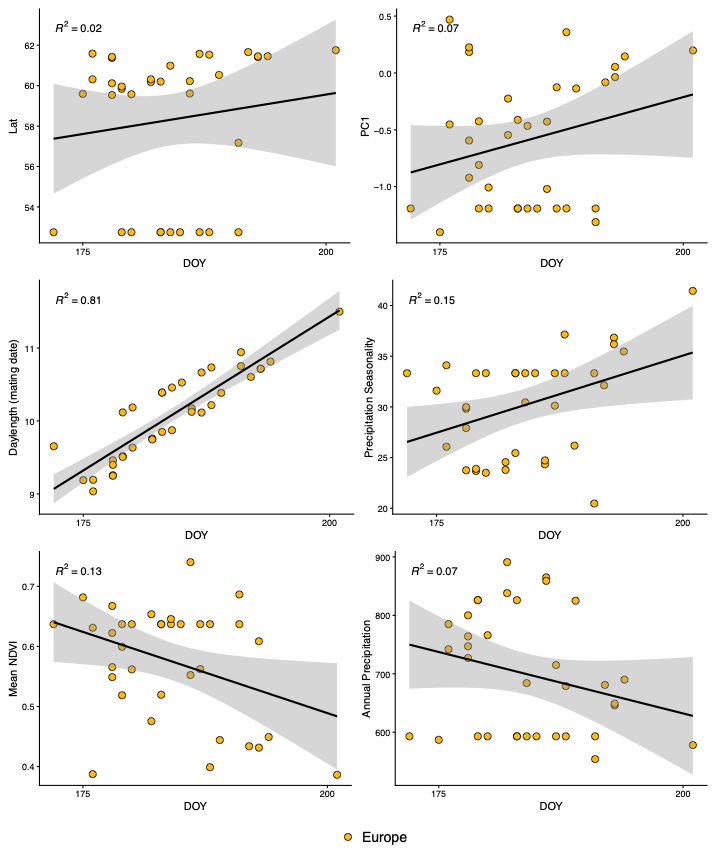

**Figure S12**. The relationship between estimated timing of birth for European wolves without wolves from Iberian peninsula and each environmental variable in our dataset, including latitude. The linear regression line is shown for each relationship and the R^2^ value. DOY is adjusted where November 1^st^ is Day 1.

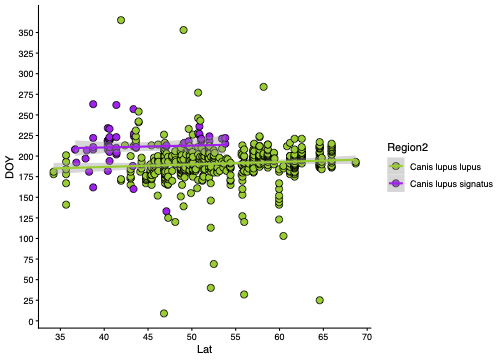

**Figure S13.** Timing of birth between 35-55N for captive wolves assigned as Eurasian wolves (*Canis lupus lupus*) and Iberian wolves (*Canis lupus signatus*). The latitudinal range of the Iberian wolf distribution is ~43.6 to ~36.2N, and captive Iberian wolves located above ~43.6N are therefore outside of the natural Iberian wolf range. DOY is adjusted where November 1^st^ is Day 1.

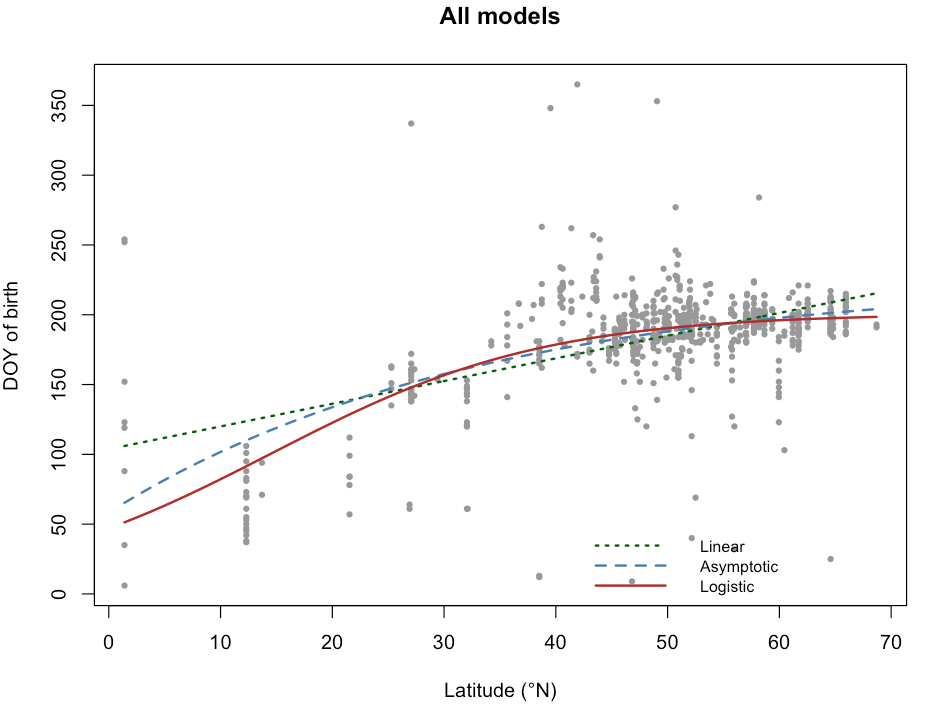

**Figure S14**. Comparison between three models (linear, asymptotic, and logistic models) for assessing the relationship between adjusted DOY and latitude for captive wolves (n=688 timings). The results of the three models is found in **Table S7**.

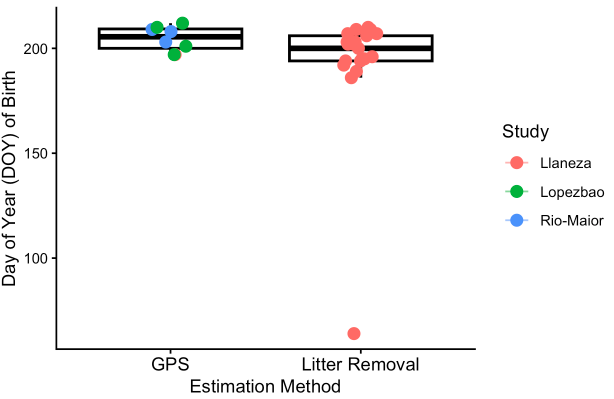

**Figure S15.** Timing of birth of our data on wild Iberian wolves (*Canis lupus signatus*). This consists of 29 timings of birth that were estimated using GPS-collared wolves and estimating age of wolf pups that were removed from dens when pups were less than 15-20 days old. Colors indicate the study that the specific data is associated with.
